# PERK-ATAD3A interaction protects mitochondrial proteins synthesis during ER stress

**DOI:** 10.1101/2022.07.24.501280

**Authors:** Daniel T. Hughes, Karinder K. Brar, Jordan L. Morris, Kelly Subramanian, Shivaani Krishna, Fei Gao, Lara-Sophie Rieder, Joshua Freeman, Heather L. Smith, Rebekkah Jukes-Jones, Jodi Nunnari, Julien Prudent, Adrian J. Butcher, Giovanna R. Mallucci

**Affiliations:** UK Dementia Research Institute and Department of Clinical Neurosciences, University of Cambridge, Cambridge Biomedical Campus, Cambridge CB2 OAH, UK; Medical Research Council Mitochondrial Biology Unit, University of Cambridge, Hills Road, Cambridge CB2 0XY, UK; Department of Molecular and Cellular Biology, University of California, Davis, Davis CA, USA; Altos Labs, Bay Area Institute of Science, Redwood Shores, CA, USA; Altos Labs, Cambridge Institute of Science, Granta Park, Cambridge CB21 6GP, UK; The Leicester van Geest MultiOmics Facility, Hodgkin Building, Lancaster Road, LE1 9HN, UK

## Abstract

Widespread repression of protein synthesis rates is a key feature of Endoplasmic Reticulum (ER) stress, mediated by the ER sensor kinase PERK. While select transcripts escape this repression, global translational down-regulation impacts crucial protein levels in all cellular compartments, beyond the ER. How the cell manages this paradox is unclear. PERK has a unique cytoplasmic loop within its kinase domain that binds PERK’s target, eIF2α. We identified the mitochondrial protein, ATAD3A, as a new interactor of the loop, binding to a highly conserved region within it. During ER stress, increased interaction between ATAD3A and PERK attenuates PERK signalling to eIF2α, removing the translational block on several mitochondrial proteins. This occurs at novel context-dependent, mitochondria-ER contact sites. The interaction provides a previously unknown mechanism for fine-tuning translational repression at a local level, mitigating the impact of ER stress on mitochondria. Further, it represents a new target for selective modulation of PERK-eIF2α signalling in diseases from cancer to neurodegeneration.

**Graphical Abstract:** 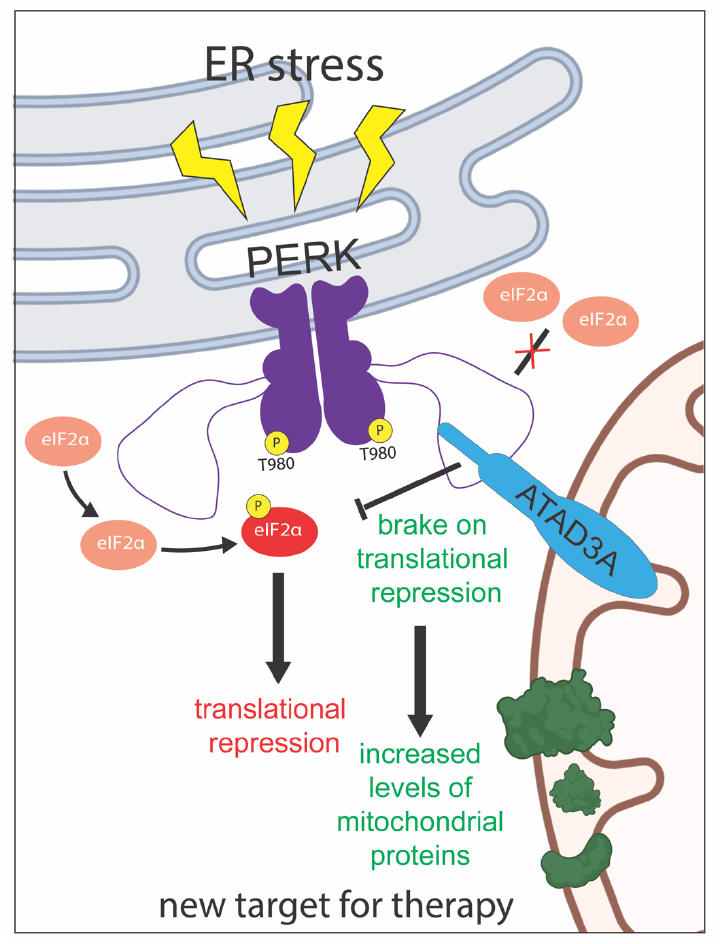

## Main text

ER stress, caused by the misfolding of proteins within the ER lumen, is detected by three transmembrane sensor/effector proteins, Inositol-Requiring Enzyme (IRE1), Activating Transcription Factor 6 (ATF6) and PKR-like Endoplasmic Reticulum Kinase (PERK), which together orchestrate the unfolded protein response (UPR) (*1*). IRE1 and ATF6 coordinate a largely transcriptional response resulting in an increase in ER chaperones and ER folding capacity (*2, 3*). In contrast, activated PERK - also a mediator of the related integrated stress response (ISR) - phosphorylates the translation initiation factor, eIF2, on its alpha subunit (*4*) resulting in the rapid reduction in global protein synthesis rates (*5, 6*). To date, the only known exceptions to global translational repression are a group of transcripts which are selectively translated during ISR/ER stress including activating transcription factor 4 (ATF4), C/EBP homologous protein (CHOP) and growth arrest and DNA damage-inducible protein 34kDa (GADD34) (*5, 7*). While there is a clear logic to translational repression within the ER whilst misfolded proteins are refolded or degraded during ER stress, the universality of this downregulation at a cellular level is paradoxical. Why should all protein synthesis – except ATF4, CHOP and related proteins - be repressed in organelles or compartments unaffected by the stress? Is there in fact a cellular mechanism for local evasion from translational repression?

PERK engages with eIF2α through an evolutionarily conserved large kinase insert loop, unique amongst protein kinases, located between the N and C-lobe of its cytoplasmic kinase domain (aa650-903) (*8*). Deletion of the loop eliminates eIF2α binding and its subsequent phosphorylation at serine 51. Phosphorylation of a single threonine residue within the loop (T799) reduces eIF2α binding, down-regulating PERK signalling (*9*), underlining the potential for regulation of the PERK-mediated response to ER stress, via its cytoplasmic face. To explore this, we performed mass spectrometry analysis to look for new PERK binding partners relevant to this phenomenon (Fig 1a, Figure S1a). This identified PERK’s luminal binding partner, GRP78 (BiP), its substrate, eIF2α, but also a new interactor, the mitochondrial protein ATAD3A, (Fig 1a), both in cells (Fig S1b) and *in vitro* (Fig 1b). ATAD3A is a nuclear encoded AAA+ domain ATPase with many reported roles, including nucleoid and inner mitochondrial membrane organisation (*10, 11*). ATAD3A spans both the mitochondrial outer and inner membranes with its C-terminal ATPase domain and N-terminal domain localized to the matrix and cytosol, respectively (*10, 12, 13*).

**Figure 1.**
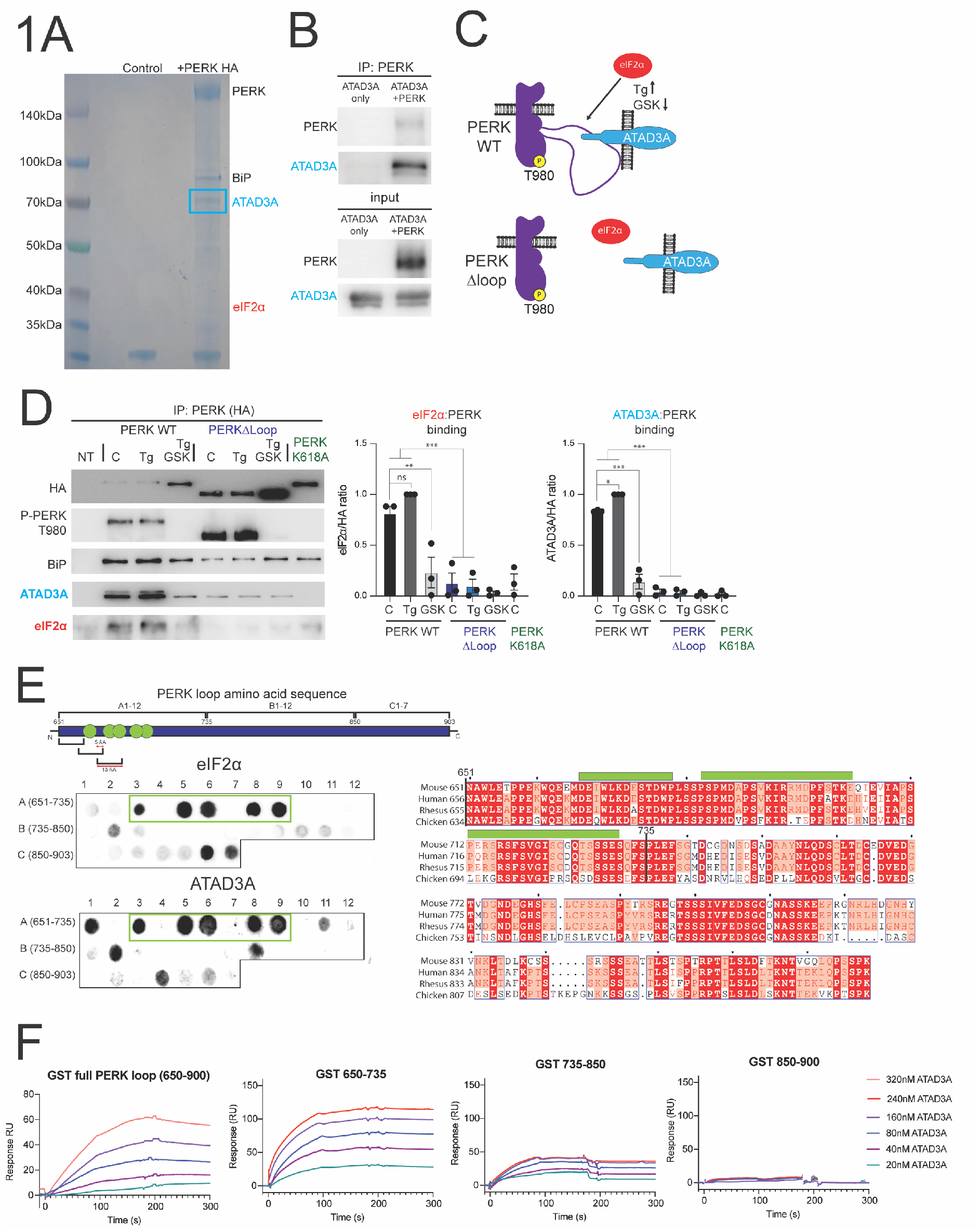
ATAD3A is a novel PERK interactor and binds to PERK’s kinase insert loop in an activity dependent manner. A) Coomassie stained SDS PAGE showing anti HA immunoprecipitation from HEK 293 cells transfected with PERK or untransfected (control). B) Association of purified PERK and purified ATAD3A *in vitro*. C) Schematic representation of PERK showing the role of the kinase insert loop in coordinating eIF2α or ATAD3A. D) Representative western blot images of anti-HA immunoprecipitates probed with antibodies to HA, PERK-P T980, BiP, ATAD3A and eIF2α, HEK 293 cells were transfected with HA tagged wild type PERK, PERK Δloop or PERK K618A then treated with DMSO (C), thapsigargin (Tg, 500nM) or thapsigargin (500nM) and GSK2606414 (10µM). Graphs show the mean ±SEM of ratios between eIF2α and HA ratio (left) ATAD3A and HA (right). Significance determined by one-way ANOVA followed by Tukey’s t test. eIF2α binding: WT C vs WT GSK (p=0.003), WT C vs Δ C (p=0.0006) and WT Tg and Δ Tg (p=<0.0001). ATAD3A binding WT C vs WT Tg (p=0.0282), WT C vs WT GSK (p=<0.0001) WT C vs Δ C (p=<0.0001) and WT Tg vs Δ Tg (p=<0.0001). n=3. E) (Top), schematic representation shows 13aa peptides derived from the kinase insert loop fractionated in to 13aa peptides which overlap by 5 aa, areas of high binding are shown (green dots) membranes were incubated with 10µg/ml of purified eIF2α or ATAD3A, each experiment was performed at least twice with independent protein preparations. (right), Schematic showing sequence alignment of PERK loop from mouse, human, rhesus macaque and chicken areas of high sequence homology are indicated. F) Examples of BIAcore T200 sensorgrams. GST fusion proteins were captured by direct immobilisation onto individual flow cells of a series S CM5 sensor chip. ATAD3A protein was diluted as indicated and perfused at 100µl/min for 3 minutes.

The ATAD3A-PERK interaction occurred selectively during PERK activation. ATAD3A-PERK binding was promoted by the ER stressor thapsigargin, and reduced by the PERK inhibitor GSK2606414 (Fig 1c, 1d, S1c, d, e). Neither ATAD3A nor eIF2α bound the PERK K618A kinase-dead mutant (Fig 1d, S1c). In addition, deletion of the kinase insert loop abolished binding of both eIF2α and ATAD3A in all conditions (Fig 1d). A similar profile of ATAD3A and eIF2α binding to PERK was confirmed in a second (murine) cell line (Fig S1d, e).

eIF2α binds directly to the insert loop (*8*), however, the precise motif/s that stabilises the interaction are unknown. To define these, we used a peptide array consisting of 13aa peptides, staggered to overlap by 5aas, covering residues aa650 to aa903 of PERK (the entire loop) incubated with purified eIF2α or purified ATAD3A (Fig 1e S1f). This gave rise to two very similar (especially at the proximal end), but distinct, binding profiles for the respective proteins. eIF2α and ATAD3A bound to peptides located at the proximal end aa682-732, a region highly conserved across species (Fig 1e), suggesting a conserved biological interface. Both bound less consistently to a second region at the distal end, aa885-903, where whilst being in the same region, the binding sites are different for the two proteins (Fig 1e). Neither GFP or GST bound to any of the array peptides, indicating that the PERK and ATAD3A interactions were specific (Fig S1g). Surface plasmon resonance (SPR) assays confirmed the ATAD3A-PERK loop binding pattern (Fig 1f). As expected, full-length and proximal loop fusion proteins bound ATAD3A in a concentration-dependent manner (Fig 1f), while mid loop and distal loop fusions had reduced or absent ATAD3A binding, respectively.

We next tested the influence of ATAD3A on PERK signalling (Fig 2a, b). Silencing ATAD3A by RNAi increased eIF2α-P levels after 30 min of thapsigargin treatment without affecting PERK phosphorylation (Fig 2b); in contrast, ATAD3A overexpression resulted in thapsigargin-treated eIF2α-P levels lower than control levels (Fig 2b), with parallel changes in ATF4, GADD34 protein and CHOP mRNA levels (Fig 2b). Thus, ATAD3A-PERK interactions attenuate PERK signalling. Recent studies support protrusion of the ATAD3A N-terminus from the outer mitochondrial membrane (OMM) (*13, 14*) into the cytosol. Electron microscopy immunogold labelling with a specific ATAD3A N-terminus antibody detected electron dense spots outside the OMM (Fig S2a), consistent with previous reports (*13*). Further, expression of mutant ATAD3A lacking the cytosolic N-terminal domain (ATAD3A Δ1-240), failed to modulate the thapsigargin-induced eIF2α-P increase (Fig 2c) or co-precipitate with PERK (Fig 2d), despite being correctly localised to mitochondria (Fig S2b). Critically, the effects of ATAD3A are specific to PERK activation and do not affect eIF2α-P signalling via other branches of the ISR, nor the other UPR branches, nor do they impact the mitochondrial-ISR. Thus, knockdown or overexpression of ATAD3A, did not affect increased eIF2α-P levels induced with the generic ISR stressor sodium arsenite (Fig S2c); nor levels of XBP1 splicing during ER stress (Fig S2d); nor the mitochondrial stress response, induced with FCCP (Fig S2e). Thus, the data support a mechanism where the N-terminus of ATAD3A localised to the outside of the mitochondria binds directly to the cytoplasmic kinase insert loop of PERK reducing its ability to bind and phosphorylate eIF2α, attenuating signalling through this pathway.

**Figure 2.**
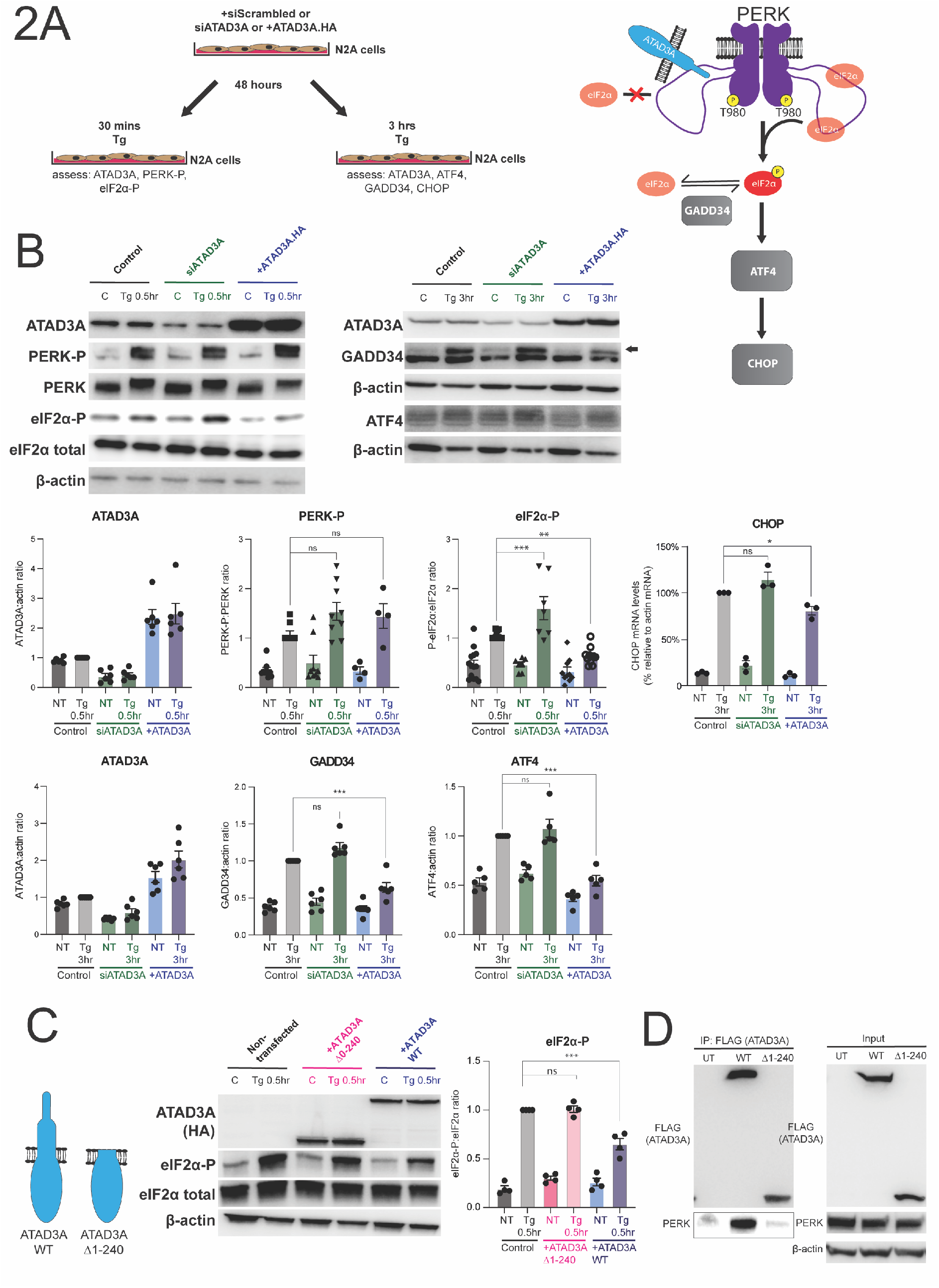
ATAD3A PERK interaction reduces PERK signalling. A) Schematic showing experimental timeline for ATAD3A knock-down/overexpression and thapsigargin treatment. Schematic showing effects of ATAD3A and mutant ATAD3A on PERK signalling. B) Western blot analysis of N2A cells transfected with scrambled siRNA, ATAD3A siRNA or HA tagged wild type ATAD3A treated with thapsigargin (500nM) or DMSO (C) for 30 mins or 3 hours. Graphs show mean ±SEM of densitometry analysis. Significance determined by one-way ANOVA followed by Tukey’s t test. For eIF2α-P: Control Tg vs siATAD3A Tg (p=0.001) Control Tg vs +ATAD3A Tg (p=0.0081). For CHOP levels: Control Tg vs +ATAD3A Tg (p=0.0484). For GADD34: Control Tg vs +ATAD3A Tg (p=<0.0001). For ATF4: Control Tg vs +ATAD3A Tg (p=0.0001). For ATAD3A blots n=6, for all conditions, PERK-P:PERK n= 9 for control cells, n=9 for siATAD3A cells and n=4 for ATAD3A HA cells and for eIF2α-P n= 14 for control cells, n=7 for siATAD3A cells and n=10 for ATAD3A HA cells, in 3 hour blots n=6 for all, ATF4 n=5. CHOP graph shows qPCR analysed levels of CHOP transcript (n=3). C) Western blot analysis of N2A cells transfected with ATAD3A WT or ATAD3A Δ1-240 and treated with thapsigargin or DMSO for 30 mins. Significance determined by one- way ANOVA followed by Tukey’s t test. For eIF2α-P: Control Tg vs +ATAD3A Tg (p=<0.0001). n=4. D) Immunoprecipitation of ATAD3A WT and ATAD3A Δ1-240 (n=3).

While PERK and ATAD3A are localised to ER and mitochondria, respectively, PERK (*15*) and ATAD3A (*10, 12*) have also each been found at mitochondria-ER contact sites (MERCS). We therefore asked whether they interact and colocalise at this specific cellular subcompartment. We examined their localization under physiological expression conditions, using U2OS cells engineered with an HA-tag on endogenous PERK (Fig S3a), and indeed observed significant colocalisation of PERK and ATAD3A (Fig 3a, upper panels). Critically, this colocalisation increased under conditions of ER stress (Fig 3a, middle panels) and was reduced by PERK inhibition (Fig 3a, lower panels) and (Fig 3b), consistent with our biochemical data (Figs 1 and 2). These points of co-localisation are likely to occur at MERCS as over-expression of PERK and ATAD3A led to increased MERCS formation, as assessed by Transmission Electron Microscopy (TEM) (Fig S3b). In contrast, expression of PERK and ATAD3A mutants that abolished their interaction, did not increase MERCS (Fig S3b). These data suggest the existence of a direct PERK-ATAD3A membrane contact site. We further quantified MERCS using TEM and found that they significantly increased during ER stress induced with thapsigargin treatment, which was prevented by pharmacological PERK inhibition (Fig 3c). Knockdown of ATAD3A (Fig S3c) similarly prevented the thapsigargin-induced increase in contacts (Fig 3d). Labelling the ER and mitochondria independently of PERK and ATAD3A further confirmed stress-dependent MERCS formation, that was inhibited by pharmacological PERK inhibition, or by siRNA of PERK and ATAD3A, respectively (Fig S4a, b and c). However, PERK inhibition or ATAD3A silencing did not affect MERCS at steady state conditions (Fig 3d, S4b). Thus, both PERK and ATAD3A are required to increase the formation of MERCS specifically in the context of ER stress, but are not necessary for the maintenance of these contacts under non-stressed conditions. This interaction therefore represents a new form of context-specific mitochondria-ER tethering.

**Figure 3.**
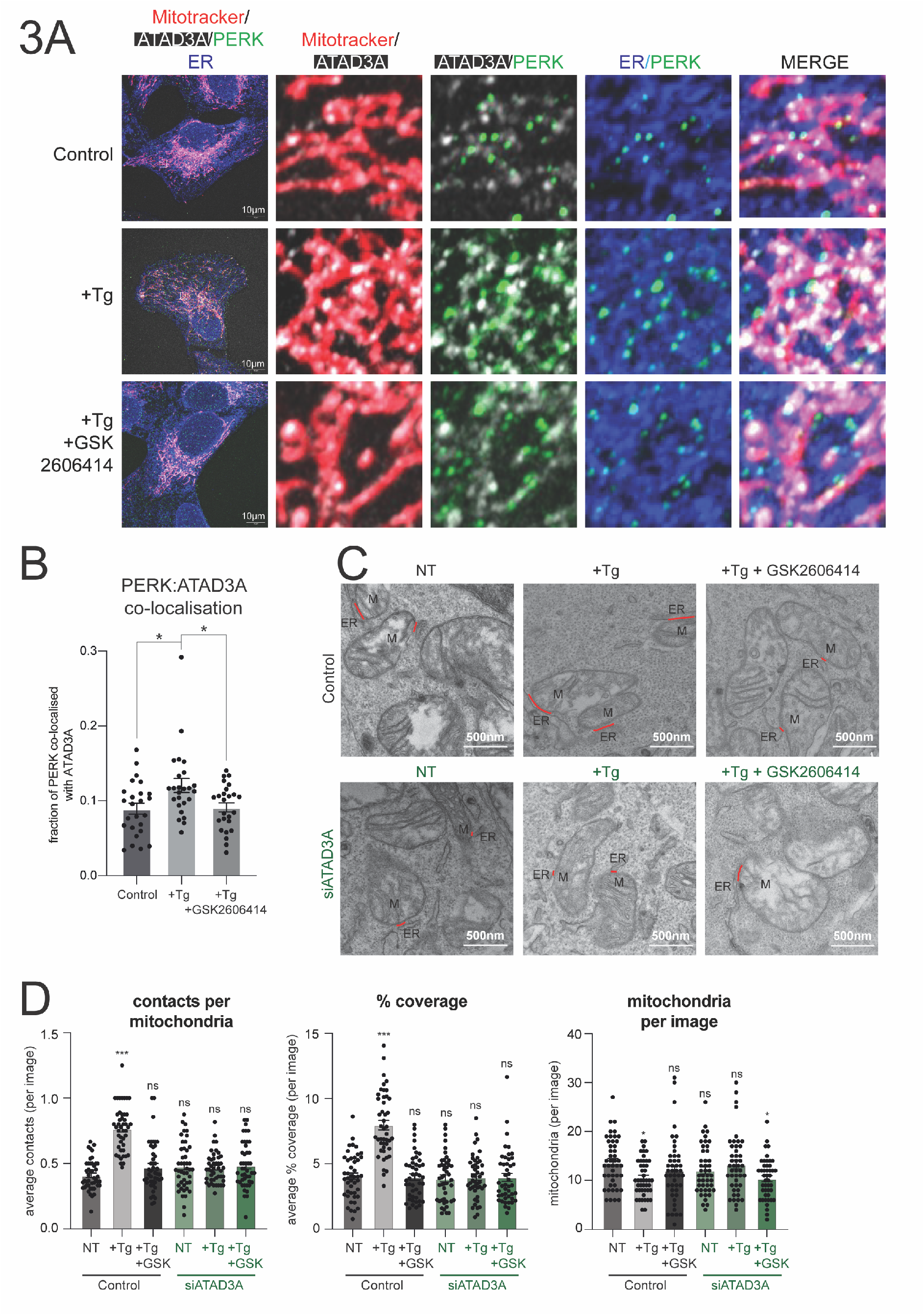
PERK and ATAD3 co-localise at juxtapositions between ER and mitochondria. A) Representative IF images of U2OS PERK HA cells treated with thapsigargin (500nM), thapsigargin and GSK2606414 (10µM) or DMSO (control) for 1 hour. Mitotracker (white), ATAD3A (pink), HA (green) and calnexin (blue) are shown in various merges. B) Graph showing levels of PERK co-localised with ATAD3A in each condition. Graph shows the mean ±SEM for each condition. Statistical significance assessed using one-way ANOVA followed by Tukey’s t test. For Control vs +Tg (p=0.016) and for +Tg vs +Tg +GSK2606414 (p=0.0246). n=25 cells from 3 separate experiments. C) Electron microscopy analysis of N2A cells treated with thapsigargin (500nM), thapsigargin and GSK2606414 (10µM) or DMSO (control) for 1 hour after being transfected with scrambled siRNA (control) or siRNA targeted to ATAD3A. Electron micrograph from control group showing (top) 10,000x magnification image used for analysis and (bottom two) zoomed image of MERCS (red line). D) Graphs showing average number of individual MERCS (per image), % coverage of mitochondria by ER (per image) and number of mitochondria (per image). Significance determined by one-way ANOVA followed by Tukey’s t test. For contacts per mitochondria: Control NT vs Control Tg (p=<0.0001). For % coverage: Control NT vs Control Tg (p=<0.0001). For mitochondria per image: Control NT vs Control Tg (p=0.0435) and Control NT vis siATAD3A GSK (p=0.0288). n=45 total images for each condition from 3 separate experiments.

Given the effects of ATAD3A on eIF2α-P levels and downstream signalling (Figs 1 and 2), our data predicts that during ER stress, the ATAD3A-PERK interaction will attenuate translational repression. To test this, we examined the proteome of thapsigargin-stressed cells in the presence or absence of ATAD3A (Tables S1, S2). Quantitative mass spectrometry and cellular compartment analysis showed that knockdown of ATAD3A selectively exacerbated the translational repression of mitochondrial proteins observed during ER stress, causing a further reduction of 27% (83 reduced in siATAD3A thapsigargin vs 45 thapsigargin alone), with no additional repression in levels of exosomal, nuclear and cytoskeletal proteins (Fig 4a). Strikingly, this was accompanied by an equivalent level of ‘de-repression’ in ER proteins (Fig 4a) suggesting that selective competitive pressure exists on levels of protein synthesis within subcellular compartments during ER stress. Further analysis showed that during ER stress, of the 20 most down-regulated mitochondrial proteins observed in the absence of ATAD3A (Fig 4b, c; Table S3), only 5 showed a marked reduction in the presence of ATAD3A, with lesser reduction of others (Fig 4b, compare middle and bottom panels). These data support the conclusion that the ATAD3A-PERK interaction plays a direct role in attenuating translational repression of a significant subset of mitochondrial proteins during ER stress. The proteins ‘rescued’ by ATAD3A during ER stress function in key regulatory nodes for mitochondrial function and biogenesis, suggesting that the PERK-ATAD3A interaction may serve to protect mitochondria from the deleterious effects of ER stress and downstream activation of the ISR. Indeed, an additional mitochondrial protective role for PERK signalling during ER stress via stress-induced mitochondrial hyperfusion has been identified (*16*). However, the mechanism underlying this protective pathway is thought to be mediated by mitochondrial lipid alterations induced by loss of the short-lived intramitochondrial phosphatidic acid transporter, PRELID1 during PERK-induced translational attenuation (*17*). In agreement, under ER stress conditions we also identified PRELID1 as a mitochondrial protein significantly down regulated; however, we observed this in both control and ATAD3A deficient cells, indicating that there are multiple PERK-dependent pathways that impinge on mitochondrial function. It is unclear at present how the translation of these specific proteins is selectively protected by the ATAD3A-PERK axis. Interestingly, many cytosolic mRNAs encoding mitochondrial proteins are translated near the OMM (*18*). Further, several OMM mRNA binding proteins (OMRBP) have been identified, of which, SYNJ2BP, which also participates in a mitochondria-ER tethering complex (*19*), has been shown to facilitate the translation and import of a specific subset of local mitochondrial proteins following recovery from translational inhibition via puromycin treatment. Although there is no apparent overlap between the mitochondrial proteins affected by ATAD3A and SYNJ2BP, they share the ability to participate in the regulation of MERCS and act at the surface of mitochondria to modulate the mitochondrial proteome via regulation of the translatome. However, the principle of local escape from global translational repression, mediated by a novel protein-protein interaction at the interface between two organelles, is a key new finding with implications for mitigation of the wider effects of ER stress on other organelles, specifically mitochondria in this case.

**Figure 4.**
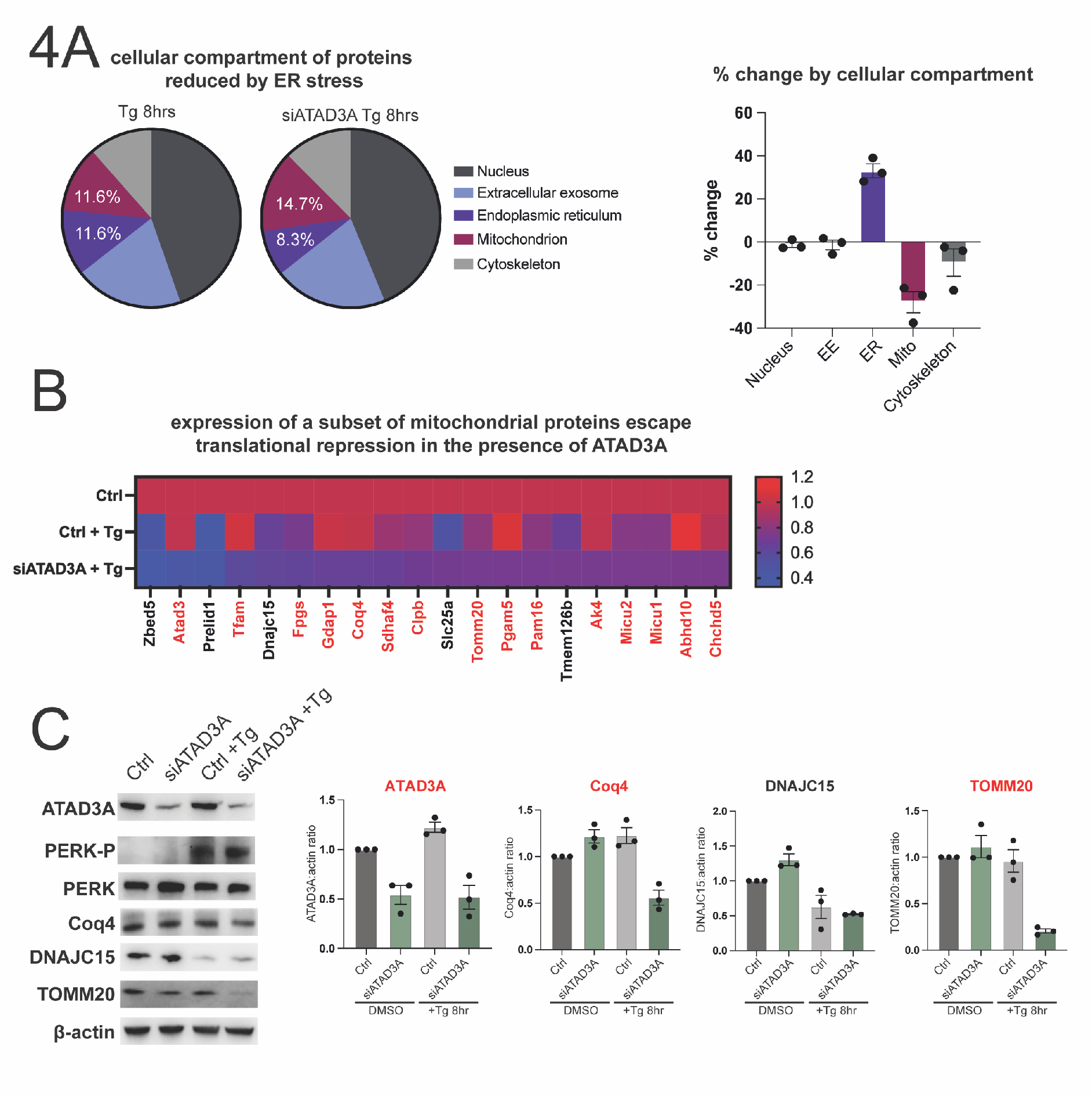
ATAD3A PERK interaction is involved in local changes in protein synthesis. A) DAVID cellular compartment analysis for conditions scrambled siRNA + thapsigargin 8 hours and siATAD3A + thapsigargin 8 hours (compared to scrambled DMSO control). Pie charts and graph showing change in proteins reduced by 20% or more by 8 hours thapsigargin treatment by cellular compartment, graph compares average of scrambled siRNA thapsigargin treated (n=2) vs siATAD3A thapsigargin treated (n=3) N2A cells. B) Heatmap showing detection of the top 20 reduced mitochondria associated proteins from siATAD3A + thapsigargin 8 condition vs all other conditions. Proteins labelled in black are most significantly reduced in both control and siATAD3A cells treated with thapsigargin (Tg). C) Representative western blot images shown to validate the mass spectrometry analysis (n=3).

In conclusion, our study uncovers a new non-canonical mechanism for the regulation of PERK signalling output in a context-dependent manner. The mechanism proposed here would result in maintenance of translation rates local to mitochondria during ER stress and a potential continuous supply of newly synthesised protein essential for proper function. Sequestration of ATAD3A in disease, as occurs in AD and HD models (*13, 20*) and FUS (*21*), could provide a basis for dysregulated PERK signalling seen in many neurodegenerative diseases (*22-24*). This mode of modulation is specific to the PERK signalling arm of the ISR due to the unique cytoplasmic insert loop present in PERK’s kinase domain. That eIF2α binds directly to the kinase insert loop indicates a mode of substrate recruitment which sets PERK aside from other kinases, including other eIF2α kinases, permitting additional modes of regulation. PERK-ATAD3A interactions further provide a new target for the selective disruption of dysregulated PERK signalling in diseases from cancer to neurodegeneration.

## Acknowledgements

We thank the Cambridge Centre for Proteomics for carrying out TMT mass spectrometry and the electron microscopy facility at the Jeffrey Cheah Biomedical Centre for carrying out TEM.

## Author contributions

DTH, KKB, JLM, KS, SK, FG, LR, JF, HLS, RJJ conducted experiments. DTH, JP, JN, contributed to study design. AJB and GRM designed the study and supervised the work. All authors contributed to writing the paper.

## Data and materials availability

All data needed to evaluate the conclusions in the paper are present in the paper or the Supplementary Materials.

