## Supplementary Materials for "PERK-ATAD3A interaction protects mitochondrial proteins synthesis during ER stress"

This file includes:
Materials and Methods
Supplementary Figures and Figure legends
Supplementary Table legends

### Materials and methods

#### Cloning

Mutant murine PERK constructs were made using PERK.WT.9E10.pcDNA3 (a gift from D. Ron, University of Cambridge) as a template, Δloop construct was amplified from a purchased cDNA from Genecopoeia. A HA tag was appended to the C terminus of WT, Δloop and K618A PERK by polymerase chain reaction using the following primers: PERK HA forward GATAGCTAGCGCCACCATGGAGCGCGCCACCCGGCCC and reverse GATACTCGAGCTAAGCGTAATCTGGAACATCGTATGGGTAGTTGCCAGGCAGTGGGCTGTACG. All constructs were verified by DNA sequencing after restriction cloning into pCDNA5 flip recombination target (FRT). ATAD3A was amplified by PCR from N2A cells, using forward primer GATCAAGCTTATGTCGTGGCTCTTCGGCATCAAGG and reverse primer GATCGCGGCCGCTCAACAACTGAGGAGTGAAGGATGTGGCG the resulting PCR product was subcloned into pcDNA5frt. ATAD3A Δ1-240 FLAG was made using the following primers, forward ACTTAAGCTTATGTCGTGGCTCTT and reverse CATGGCGGCCGCCTATTTATCGTCATCGTCCTTATAGTCCACAAATGCACGGAATCCTTCCAC and was subcloned into pcDNA5frt. The C-terminus of ATAD3A was modified such that an HA epitope tag or a twin-strep-tag were fused to the final coding amino acid the following primers were used; ATAD3A HA reverse GATAGCGGCCGCCTAAGCGTAATCTGGAACATCGTATGGGTAACAACT GAGGAGTGAAGGATGTGGCG, ATAD3A twin-strep reverse GATACTCGAGCTACTTCTCGAACTGGGGGTGGCTCCAGGCGGAGCTGCCTCCGCTGCCGCCGCCGC TGCCGCCTCCTTTTTCAAATTGAGGATGAGACCATGCAGAACAACTGAGGAGTGAAGGATGTGGCG. GST fusions of the PERK kinase insert loop were generated by PCR using the following primers;

650 forward GATCGGATCCTTCAACGCCTGGCTGGAAACCCCACC, 902 reverse GATCGAATTCTCA TGAGCAGCGCCGGTTCATCCAGTC, 735 forward GATC GGATCCTCTCCCCTGGAGTTCTCAGGGACAG, 735 reverse GATCGAATTCTCAAGAGAACTGGCTCTCCGATGAGC, 847 forward GATCGGATCC GAAGCCACCACCTTGTCTACCTCC, 847 reverse GATCGAATTCTCATTCTGAAGAAGACCTGCTGCTGGAGC. The resulting products were subcloned into pGEX-2T.

**Protein expression and purification**

For protein expression in *E. coli*, BL21 C+ RIL containing constructs in pGEX-2T were grown in LB until OD@600nm = 0.6. Cultures were cooled to 16°C and protein production was initiated by addition of IPTG to a final concentration of 0.2mM, cultures were grown for a further 4 hours before harvesting.

Cell pellets representing 1 litre of induced culture were re-suspended in 20ml PBS containing 1mM DTT with protease inhibitors (Complete, Roche) and lysed using a probe sonicator with 4 x 30 seconds at 50% power. Following centrifugation @ 20,000 x g for 30 minutes at 4°C, the resulting supernatants were loaded onto 1.5ml glutathione sepharose (GE healthcare), washed with 50 volumes of PBS and eluted in 5 volumes of PBS containing 20mM reduced glutathione. Purified GST fusions were stored at -80°C.

PERK twin-strep or ATAD3A twin-strep were expressed in EXPI293 cells. For purification, cell pellets representing 1 litre of culture were re-suspended in 20 ml TE containing protease inhibitors and lysed by sonication. Lysates were diluted to 100ml and centrifuged at 35,000 x g for 1 hour at 4°C. The resulting pellet was resuspended in 20 ml buffer containing 50mM Tris pH 8.0, 150mM NaCl, 1% n-Dodecyl-β-D-Maltoside (DDM) (Thermo Scientific). Following centrifugation at 20,000 x g the resulting supernatant was diluted 1:1 with 50mM Tris pH 8.0, 150mM NaCl, and applied to a 1.5ml Streptactin-XT superflow (IBA life sciences) column. The column was washed with 100 volumes of buffer containing 0.1% DDM and eluted in 10 volumes of buffer containing 0.1% DDM and 50mM biotin. Good fractions were pooled, concentrated and stored at 4°C.

**PERK:ATAD3A complex formation *in-vitro* and co-immunoprecipitation**

Purified PERK (2 µg) and ATAD3A (8 µg) were incubated in 50 µL binding buffer [PBS, 0.5 mM ATP, 1 mM $MgCl_2$, 0.02% Dodecyl-β-D-maltopyranoside (DDM, Sigma-Aldrich)] overnight, rotating at 4°C. Samples were diluted 1:10 in binding buffer and incubated with 1 mg Protein G Dynabeads (Invitrogen) conjugated to 2 µg anti-PERK antibody (SantaCruz Biotechnologies, #9477) for one hour at room temperature. Beads were washed five times with PBS, 0.02% DDM, 0.01% Triton-X 100 (Sigma-Aldrich), resuspended in 50 µL 4x SDS loading buffer and boiled at 95°C for 10 minutes. Samples were analysed by SDS-PAGE and Western Blotting.

**Cell lines and transfection**

All cells were incubated at 37°C with 5% $CO_2$ in T75cm$^2$ or T175cm$^2$ flasks. HEK 293 and N2A cells were both maintained in Dulbecco's modified Eagle's medium (DMEM) (Gibco, 61965) supplemented with 10% fetal bovine serum (FBS) and 1% penicillin/streptomycin and 1X nonessential amino acids, and U2OS cells were grown in high glucose DMEM supplemented with 10% FBS and 1% penicillin/streptomycin. Cells were plated the day before transfection in normal medium in six-well plates (200,000 cells per well) or in T175cm$^2$ tissue culture flasks (5 million cells per flask). Cells in six-well plates were transfected with 500 ng of plasmid DNA or 10/100nM as indicated of siRNA using Lipofectamine 3000 (using manufacturer's standard protocol). Cells in T175cm$^2$ flasks were transfected with 6 µg of plasmid DNA using PEI at a ratio of 6:1 PEI:DNA. For all transfections, medium was changed after 6 hours, and samples were collected 48 hours after transfection.

For expression of ATAD3A twin-strep or PERK twin-strep, EXPI293 cells were maintained in EXPI293 expression medium in suspension cultures at 120RPM at 37°C with 8% $CO_2$. Cells were seeded at 2 x $10^6$ / ml and transfected with 1µg DNA / ml culture using PEI. Cells were supplemented with 1% pen/strep after 24 hours and harvested 72 hours post-transfection.

**Immunoprecipitation of HA-tagged proteins**

HEK 293/N2A cells from T175cm$^2$ flasks were suspended in 10 ml of 10 mM tris/1 mM EDTA supplemented with protease and phosphatase inhibitors (cOmplete, Mini, EDTA-free, Sigma-Aldrich; PhosSTOP, Sigma-Aldrich). After sonication membranes were fractionated by centrifugation at 40,000g for 45 min at 4°C, membrane pellet was resuspended in phosphate-buffered saline (PBS) containing 1% dodecyl-β-d-maltopyranoside (DDM) (Sigma-Aldrich) and incubated on ice for 20 min. Insoluble material was removed by centrifugation at 20,000g for

20 min at 4°C. Samples were diluted 1:1 in PBS for 0.5% DDM final concentration. 50 µl (for SafeStain gel) or 10 µl (for western blotting) of Anti-HA Affinity Matrix (Sigma-Aldrich) was added, and samples were incubated overnight at 4°C on a roller. HA beads were pelleted at 1000g and washed three times in PBS containing 0.1% DDM, in which the NaCl concentration was increased to 500 mM for a final wash to remove ionic interactions. Beads were then resuspended in 50 µl of 2X SDS loading buffer before SDS–polyacrylamide gel electrophoresis (PAGE) and staining or Western blotting. Gels containing samples for mass spectrometry were stained using SimplyBlue™ SafeStain (ThermoFisher Scientific).

**Peptide array to identify PERK:ATAD3A and eIF2α interaction sites**

Peptide arrays were generated by Genscript with peptides C-terminally coupled to a cellulose membrane via a β-alanine spacer. The peptides consisted of 13 amino acids with a 5 amino acid overlap (the N' terminal 5 amino acids overlapping with previous peptide) starting from amino acid 651 to 903 of murine PERK. Before use, the membrane was rehydrated in methanol for 5 minutes and thoroughly washed with TBS. The membrane was then blocked in TBS containing 0.1% Tween-20 (TBS-T) with 3% BSA for 2 hours at room temperature before eIF2α (Sigma SPR5232, EIF2S1 recombinant His tagged human protein) or strep-tagged ATAD3A was added to the blocking buffer at a concentration of 5µg/ml and incubated overnight at 4°C. Detection of bound protein to peptide array required sequential incubation with eIF2α (Rb-anti eIF2α XP CST #53245) or ATAD3A (Strep-MAB IBA-Lifesciences) polyclonal antibodies at 1:5000 dilution in blocking buffer for 2 hours at room temperature and incubation with Goat Anti-Rb and Goat Anti-Ms HRP-conjugated antibodies at 1:5000 dilution in blocking buffer for 1 hour at room temperature. The membrane was extensively washed with TBS-T between protein and antibody incubations. For chemiluminescent detection, a 1:1 ratio of ClarityTM Western Luminol/Enhancer Reagent (Bio-Rad) and ClarityTM Western Peroxide Reagent was added to the membrane and HRP antibodies were detected by the ChemiDoc chemiluminescent Imager. For control experiments, the above was repeated except the membrane was not incubated with eIF2α or ATAD3A protein.

**SPR binding assay**

Assays were performed on a BIAcore T200 instrument at 25°C in a running buffer of 20mM HEPES pH7.5 150mM NaCl, 1mM EDTA, 0.005% TWEEN-20. GST fusion proteins were covalently attached to the surface of CM5 dextran sensor chips according to manufacturer's instructions. Approximately equal molar quantities of each protein were applied, 1000 Resonance Units (RU) of GST alone, 1900 RU of GST 650-902, 1300RU of GST 650-735, 1400 RU of GST 735-850 and 1200 RU of GST 850-902. ATAD3A was diluted in running buffer and applied at 100µl/min for 3 min followed by a period of dissociation.

Sensorgrams were processed manually by aligning the injection start and end points and were corrected for nonspecific interactions by subtraction of the corresponding sensorgram recorded from the flow cell containing GST only.

**Cell treatments**

Cells were incubated in their normal media in the presence or absence of various compounds for the times specified before preparation of samples. The compounds were dissolved in DMSO and added directly to the culture media at the following final concentrations: thapsigargin (500 nM; Thermo Fisher Scientific), GSK2606414 (10 µM; GlaxoSmithKline), sodium meta-arsenite (500 µM; Sigma-Aldrich) and FCCP (25µM, Abcam). siRNA to ATAD3A (GACAGGACAGCACAGUAGUAA) or scrambled control (GUGGCUCUCGCGAUUGACUAA) were used at 100nM for mouse cells. For human cells PERK esiRNA (Sigma, EHU030881) and ATAD3A (Santa Cruz, sc-88047) were used according to manufacturer's protocol.

Cell pellets were scraped into 200 µl of radioimmunoprecipitation assay (RIPA) buffer containing protease and phosphatase inhibitors (cOmplete, Mini, EDTA-free, Sigma-Aldrich; PhosSTOP, Sigma-Aldrich) and kept on ice for 30 min with vortexing every 10 min. Insoluble material was removed by centrifugation at 15,000g for 20 min at 4°C, and supernatant was collected and frozen at −80°C. All protein samples were quantified using Bradford Assay (Bio-Rad).

**Western blotting**

10 to 30 µg of each sample were separated by SDS-PAGE on 10% tris/glycine gels, followed by transfer onto polyvinylidene difluoride membranes. Membranes were blocked in 5% bovine serum albumin for an hour at room temperature before adding primary antibodies and incubated overnight on a shaking platform at 4°C. Antibodies to ATAD3A (ab67992,) were purchased from Abcam. β-actin (#4970; 1:5000), eIF2α (#2103; 1:2000), eIF2α (D7D3) XP (for IP samples) (#5324; 1:5000), eIF2α-P Ser51 (#3597; 1:1000), HA (Sigma-Aldrich,

#11867423001; 1:5000), PERK (#3192; 1:2000), and PERK-P Thr980 (#3179; 1:2000) antibodies were purchased from Cell Signalling Technology and antibodies to ATF4 (#sc-200; 1:1000) and GADD34 (10449-1-AP) were purchased from Santa Cruz Biotechnology and Proteintech, respectively. Membranes were washed three times on a shaking platform with gentle agitation for 10 minutes with tris buffered saline with 0.1% tween 20 (TBS-T) before incubation with horseradish peroxidase–conjugated secondary antibodies for 1 hour at room temperature. Blots were then detected using enhanced chemiluminescence, and immunoreactivity was quantified using Fiji software.

**qPCR analysis**

RNA was extracted using the RNeasy micro kit and reversed transcribed using the Superscript IV First Strand system (Thermo). qPCR was performed using Power SYBR Green and 300 uM of the following primers: CHOP CCTGAGGAGAGAGTGTCCAG, ATGTGCGTGTGACCTCTGTT (FWD + RVS), XBP1 spliced GCTGAGTCCGCAGCAGGT, CAGGGTCCAACTTGTCCAGAAT (FWD + RVS), XBP1 total TGAAAAACAGAGTAGCAGCGCAGA, CCCAAGCGTGTTCTTAACTC (FWD + RVS) as in (*25*) and Actin AGTGTGACGTTGACATCCG, GCCAGAGCAGTAATCTCCT (FWD + RVS). Actin primers were used at 100 uM. qPCRs were run on a QuantStudio 7 Real Time PCR machine (Applied Biosystems) and analysed using QuantStudio Real Time software (Applied Biosystems).

**Generation of PERK HA U2OS line**

An HA tag was added to the C-terminus of endogenous PERK by transfecting U2OS cells with guide RNA 5' ATTGCTTGGCAAAGGGCTAT in vector px458 and double stranded repair oligonucleotide 5' CTTGAGTTCATCGGGAACAAAACATTCAAGACAGTCCAACAACTCCCATAGCCCTTTGCCAAGCAAT tacccatacgatgttccagattacgcttgaCCTTAAGTTGTGCTAGCAACCCTAATAGGTGATGCAGATAATAG CCTACT. GFP positive cells were selected and clones screened by PCR using the following primers 5' GATGGTTCAAGACATGCTCTC and 5' AAGAGACTAACAAAGAACAAAGATAGC. Positive clones were confirmed by sanger sequencing and western blotting.

**Immunofluorescence Microscopy**

For PERK:ATAD3A co-localisation cells were seeded at ~0.25x10^5 per mL in 35mm glass-bottom dishes (Mattek), and allowed to attach for 24 to 48 hours prior to mitochondrial staining with 500nM Mitotracker Deep Red (ThermoFisher; catalog no. M22426) according to manufacturer's instructions. Where indicated, cells were treated with thapsigarin (500nM) or co-treated with thapsigargin (500nM) and GSK2606414 (10uM) for 1 hour. Cells were then directly fixed in 4% paraformaldehyde in PBS pH 7.4 for 20 minutes at room temperature, protected from light. Dishes were washed twice in PBS, permeabilized in 0.1% TritonX-100 for 15 minutes, and blocked in 5% BSA PBS (blocking solution) for 1 hour at room temperature. Primary antibodies (Calnexin rabbit 1:250, HA rat 1:1000) were added in blocking solution and incubated 1 hour at room temperature or overnight at 4 C. Dishes were then rinsed 5 times in PBS and incubated with Alexa-Fluor conjugated secondary antibodies (ThermoFisher) at 1:1000 dilution in blocking solution for 45 minutes at room temperature, rinsed 5 times in PBS. A custom antibody targeted to the N-terminal domain of ATAD3A was made by Eurogentec using their speedy rabbit protocol with peptide CRGLGDRPAPKDKWSN and used in IF experiments. ATAD3A rabbit antibody was directly conjugated to Alexa Fluor 555 using Zenon Rabbit IgG labelling Kit (ThermoFisher; catalog no. Z25305). Rabbit Calnexin was purchased from Proteintech (10427-2-AP). Imaging was performed using the Airyscan LSM 980 with the 63X, 1.4 NA objective. Images were captured with Zen Blue 3.5; linear adjustments and quantification analyses were performed in FIJI (NIH). To quantify the fraction of PERK co-localized with ATAD3A, threshold was performed using max z-projections in FIJI of images taken under all the same imaging conditions for each experiment. Threshold 555- and 488-channel images were used to segment localization of PERK and ATAD3A. Area of PERK and ATAD3A were identified from segmentation and the overlap regions were determined using the ROI manager in FIJI.

For ATAD3A Δ1-240 imaging, cells were fixed in 4% paraformaldehyde (PFA) in PBS at 37°C for 15 min, then washed 3 times with PBS, followed by quenching with 50 mM ammonium chloride in PBS. After 3 washes in PBS, cells were permeabilized in 0.1% Triton X-100 in PBS for 10 min, followed by 3 washes in PBS. Cells were then blocked with 10% goat serum in PBS, followed by incubation with primary antibodies to TOMM20 (Alexa Fluor 647, ab209606), FLAG (Sigma, F3165) in 5% goat serum in PBS, for 2 hr at room temperature (RT). After 3 washes with 5% goat serum in PBS, cells were incubated with secondary antibody (mouse

488, 1:1000) for 1 hr at RT. After 3 washes in PBS, coverslips were mounted onto slides using ProLong™ Diamond Antifade (Thermo Fisher Scientific) before imaging on Leica Stellaris 8. For KDEL/TOMM20 contact counting cells were fixed in 4% PFA in PBS pH 7.4 at 37°C for 15 min. Cells were then washed and quenched by incubating cells with 50 mM ammonium chloride for 10 min at RT and then washed three times with PBS. After fixation and PFA autofluorescence quenching, cells were permeabilised in 0.1% Triton-X-100 in PBS for 10 min at RT. After three PBS washes, cells were blocked in 10% FBS in PBS for 20 min at RT. Cells were then incubated with primary antibody in 5% FBS in PBS overnight at 4°C with gentle agitation. Post incubation, primary antibody was removed by performing three 5 % FBS in PBS washes. Cells were then incubated with the appropriate Alexa fluor-conjugated secondary antibody (Invitrogen) 5% FBS in PBS for 1 hr at RT. Images were acquired using an Andor Dragonfly spinning disk confocal system equipped with a Nikon Eclipse Ti-E microscope and a Plan-Apochromat 100X/1.45 NA oil immersion objective, coupled with an Andor Zyla 4.2 PLUS sCMOS 4.2-megapixel camera. Images were acquired using appropriate lasers and with the Fusion software (Andor). 7 Z- stacks were acquired at 0.2 µm steps with the same laser intensity, exposure time and camera settings for all conditions in each independent experiment. Images were then compiled by 'max projection' and processed using Fiji (ImageJ) software. To analyse the interaction between mitochondria and ER by confocal microscopy or N- SIM, colocalization analysis was performed in Fiji. Briefly, z-stacks of cells immunostained with anti-TOMM20 and anti-KDEL antibodies were acquired and processed as previously described. Max projected images were then manually thresholded to remove background signal and colocalization between anti-TOMM20 and anti-KDEL signal was determined using the 'Coloc 2' plugin and a Mander's colocalization coefficient was calculated.

**Electron microscopy**

N2A cells grown on coverslips were washed twice in warm PBS after stated treatment, then fixed in 0.1M sodium cacodylate containing 4% paraformaldehyde and 2.5% glutaraldehyde overnight at 4°C. Cells were then transferred into 0.1M sodium cacodylate. Samples were post-fixed in 1ml of 1% osmium tetroxide; 1.5% potassium ferricyanide overnight at 4°C before washing in water. *En bloc* staining was carried out in 2% aqueous uranyl acetate overnight at 4°C to enhance final contrast. Samples were washed with water before dehydration by ethanol series (70%; 95%; 100%) and incubated for a further 5 mins in 100% ethanol. Dehydrated samples were incubated in 1ml 1:1 mixture of absolute ethanol: TAAB resin for 1 hr 30 minutes. Resin mixture was replaced with fresh pure resin for 2x 4hr and replaced with pure resin and incubated for 1 hr 30 minutes. Excess resin was removed, and the coverslips were placed onto aluminium foil. Flat-bottom BEEM capsules were filled with fresh resin and inverted onto the coverslips followed by polymerised at 60°C for at least 24h. Samples were sectioned at 50nm using a Leica EM UC7 ultramicrotome. Sections were mounted onto 200mesh Formvar/carbon copper grids. After counter-staining with lead citrate, samples were imaged on a Hitachi HT7800 TEM.

**Quantification of Mitochondrial-ER contact sites from TEM images**

EM images of N2A cells were taken at 10,000x magnification before analysis using Image J software. MERCS were defined as any area where mitochondrial membrane and ER were within 30nm (measured using Image J software). The number and perimeter of all mitochondria in an image was measured as well as the total number of contacts and the combined length of those contacts.

**Tandem mass tag proteomics**

N2A cells were plated in T175cm$^2$ flasks at 5 million cells per flask, 24 hours later they were transfected at 100nM with scrambled or ATAD3A siRNA using Lipofectamine 3000, 48 hours later they were treated with DMSO or thapsigargin (500nM) for 8 hours. Cells were then scraped into RIPA buffer before protein quantification. 200µL of lysate at 100µg/µL was sent to the Cambridge Centre for Proteomics for TMT labelling and mass spectrometry. Data was analysed using Microsoft excel and the Database for Annotation, Visualization and Integrated Discovery (DAVID).

**Statistical analysis**

Data are presented as mean ± standard error of the mean (SEM) unless otherwise specified in the legend. Statistical significance was determined using GraphPad Prism v9, using indicated tests. Statistical significance was accepted at $P \leq 0.05$. In the figure legends, "ns" denotes $P \geq 0.05$, * denotes $P \leq 0.05$, ** denotes $P \leq 0.01$, and *** denotes $P \leq 0.001$

### Supplementary Figures and Legends:

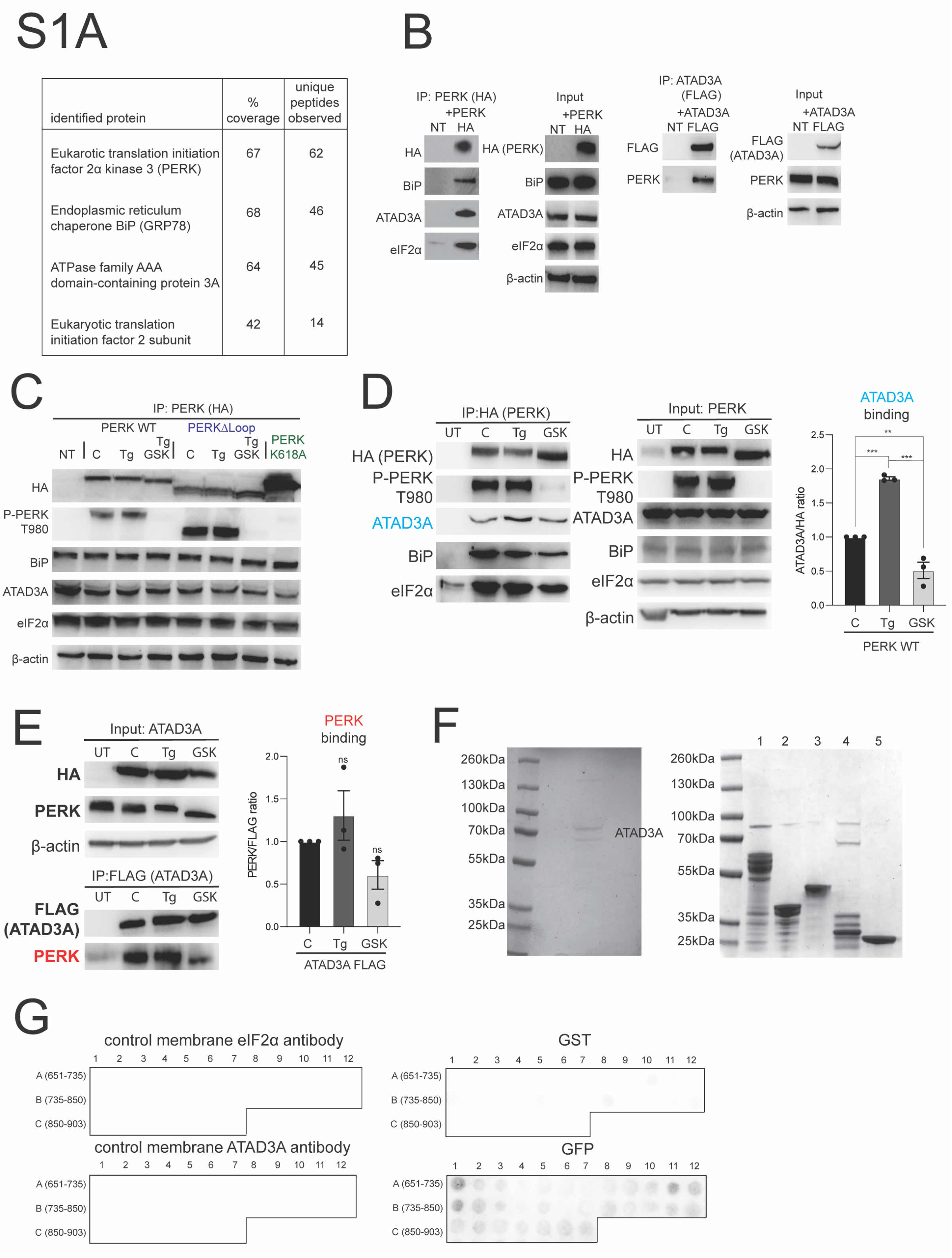


| identified protein | % coverage | unique peptides observed |
|---|---|---|
| Eukarotic translation initiation factor 2α kinase 3 (PERK) | 67 | 62 |
| Endoplasmic reticulum chaperone BiP (GRP78) | 68 | 46 |
| ATPase family AAA domain-containing protein 3A | 64 | 45 |
| Eukaryotic translation initiation factor 2 subunit | 42 | 14 |

**Supplementary Figure 1**

(A) Table showing protein identification of PERK, BiP, ATAD3A and eIF2α, including percentage coverage and number of unique peptides observed.

(B) Representative western blot images of HA, BiP, ATAD3A, eIF2α, FLAG and PERK using HA or FLAG immunoprecipitates from HEK 293 cells transfected with HA tagged wild type PERK or FLAG tagged wild type ATAD3A (NT = not transfected). n=3.

(C) Representative input blots from samples used in Fig 1d, n=3.

(D) Representative western blots of HA (PERK), PERK-P T980, ATAD3A, BiP and eIF2α, from HA-immunoprecipitated samples after transfection with HA-tagged wild type PERK expressed in N2A cells. Relevant input blots included. Graph shows mean ±SEM of ATAD3A:HA ratio. Significance determined by one-way ANOVA followed by Tukey's t test. For C vs Tg (p=0.0004), for C vs GSK (p=0.0069) and for Tg vs GSK (p=<0.0001). n=3.

(E) Representative western blots of FLAG (ATAD3A) and PERK from FLAG immunoprecipitated samples after N2A cell transfection with FLAG-tagged wild type ATAD3A. Relevant input blots included. Graphs show mean ±SEM of PERK:FLAG ratio. Statistical significance assessed using one-way ANOVA followed by Tukey's t test.

(F) Coomassie stained SDS PAGE gel showing purified ATAD3A used in peptide array and Biacore experiments. Coomassie stained SDS PAGE gel showing various GST tagged PERK loop constructs used in Biacore experiments.

(G) Representative images of control membranes for Fig 1e.

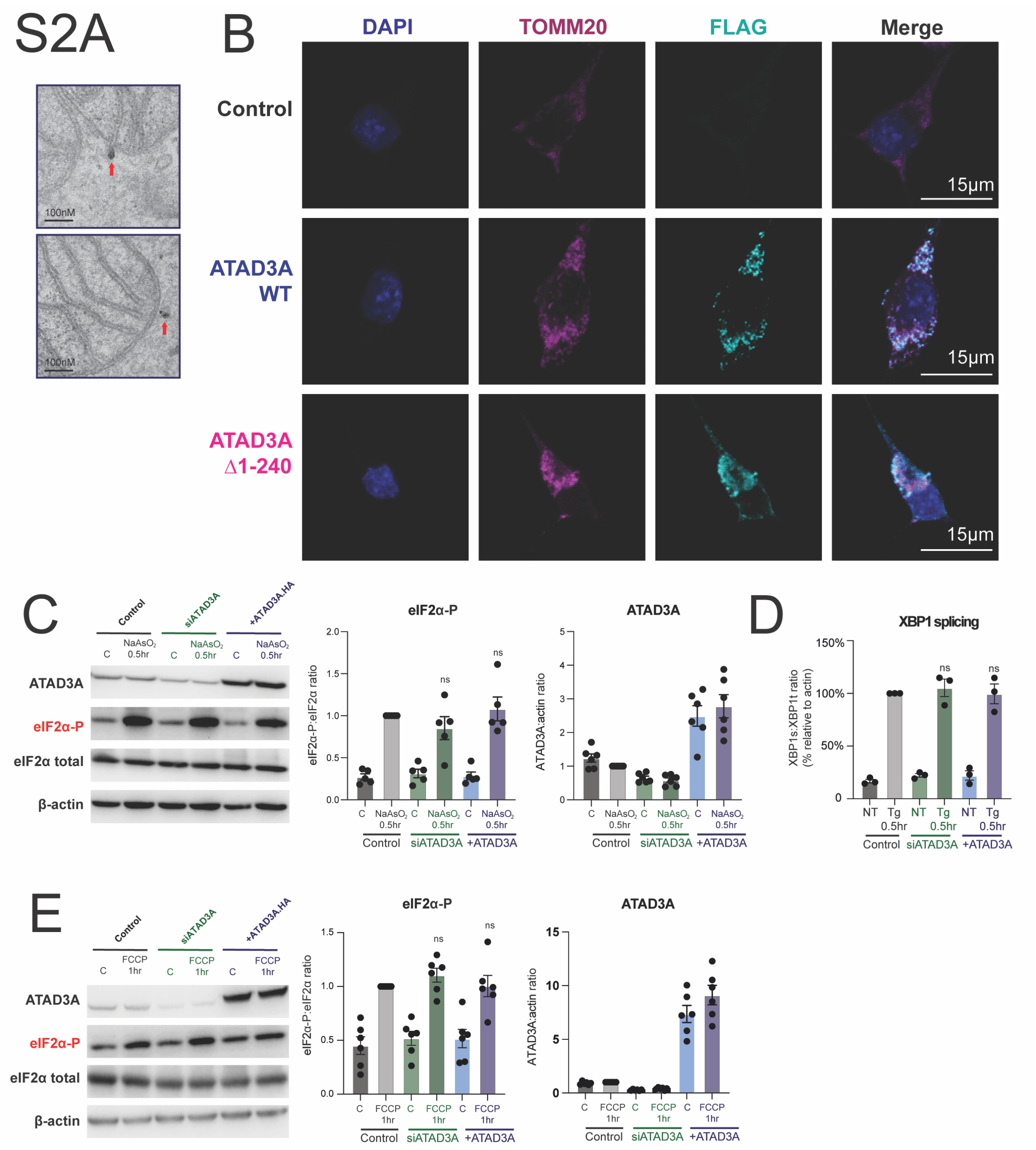
S2A
100nM
B
DAPI
TOMM20
FLAG
Merge
Control
ATAD3A WT
ATAD3A △1-240
15μm
C
Control
siATAD3A
+ATAD3A.HA
NaAsO2 0.5hr
ATAD3A
eIF2α-P
eIF2α total
β-actin
eIF2α-P
eIF2α-P:eIF2α ratio
ATAD3A
ATAD3A:actin ratio
ns
D
XBP1 splicing
XBP1s:XBP1t ratio (% relative to actin)
NT
Tg 0.5hr
siATAD3A
+ATAD3A
E
FCCP 1hr

**Supplementary Figure 2**

(A) Immunogold labelling of ATAD3A with an N-terminal targeted antibody confirms it protrudes from the mitochondria into the cytosol, n=1.

(B) Representative IF images showing DAPI, TOMM20 (alexfluor 647), FLAG and a merge from N2A cells transfected with either ATAD3A WT FLAG or ATAD3A Δ1-240 FLAG untransfected as a control.

(C) Representative western blots showing levels of HA (ATAD3A), eIF2α-P, eIF2α and actin in N2A cells transfected with siRNA to ATAD3A or ATAD3A HA and treated with Sodium Arsenite (500nM, 30 mins) or DMSO as a control. Graphs show mean ±SEM. Statistical significance assessed using one-way ANOVA followed by Tukey's t test. ns = non-significant. n=5 or 6 as displayed.

(D) qPCR analysis of XBP1 splicing in cells treated with siRNA to ATAD3A or over-expressing ATAD3A 48 hours before treatment with thapsigargin (500nM 30 mins) or DMSO control. Data presented as mean ±SEM of XBP1spliced: XBP1total ratio relative to actin control. Statistical significance assessed using one-way ANOVA followed by Tukey's t test. ns = non-significant. n=3.

(E) Representative western blots showing levels of HA (ATAD3A), eIF2α-P, eIF2α and actin in N2A cells transfected with siRNA to ATAD3A or ATAD3A HA and treated with FCCP (25µM, 1hr) or DMSO as a control. Graphs shows mean ±SEM of eIF2α-P:eIF2α ratio and HA (ATAD3A):actin ratio. Statistical significance assessed by one-way ANOVA followed by Tukey's t test. ns = non-significant. n=6

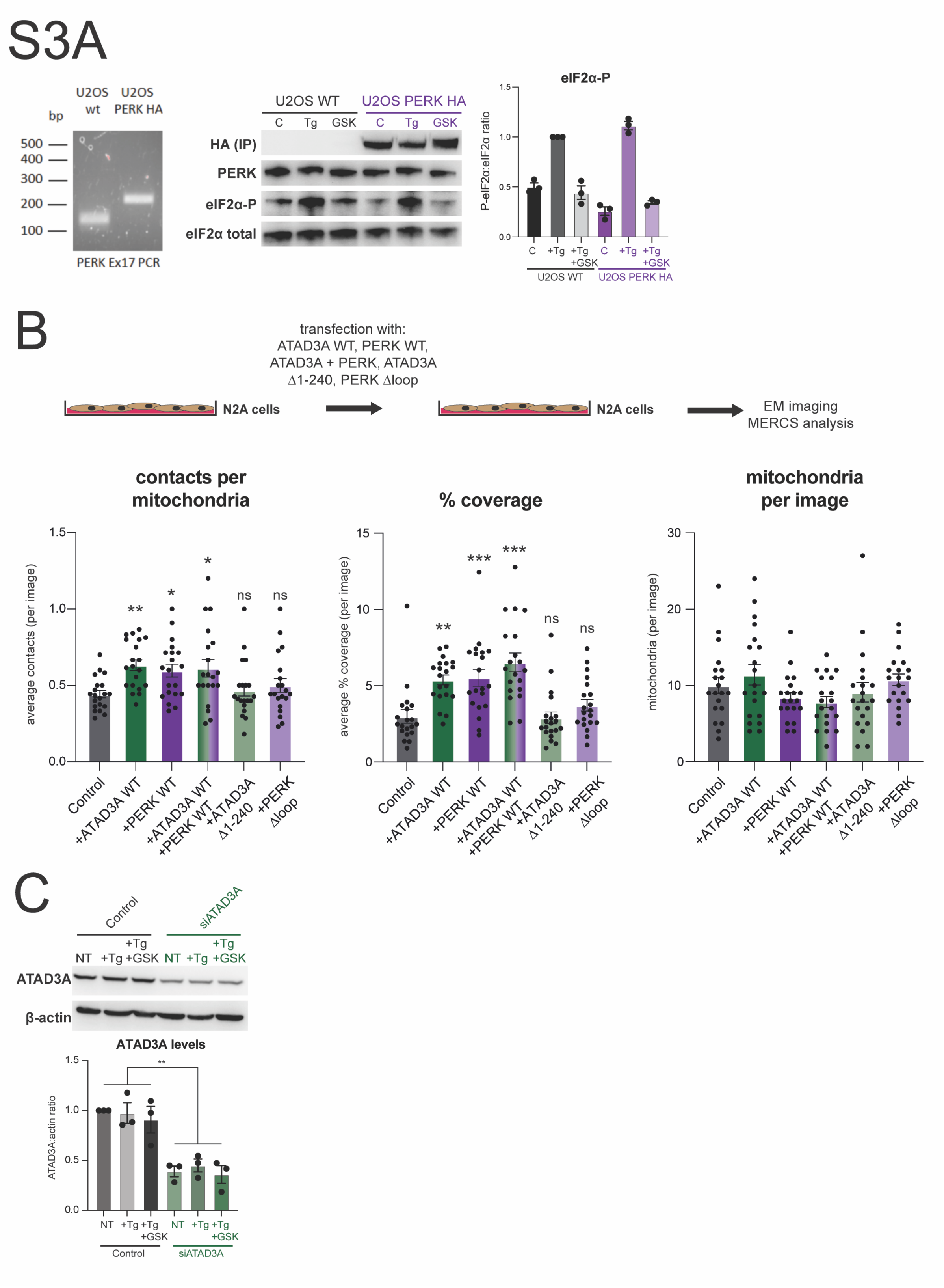

S3A
U2OS wt
U2OS PERK HA
bp
500
400
300
200
100
PERK Ex17 PCR
U2OS WT
U2OS PERK HA
C Tg GSK C Tg GSK
HA (IP)
PERK
eIF2α-P
eIF2α total
eIF2α-P
P-eIF2α:eIF2α ratio
C +Tg +Tg +GSK C +Tg +Tg +GSK
B
transfection with:
ATAD3A WT, PERK WT,
ATAD3A + PERK, ATAD3A
Δ1-240, PERK Δloop
N2A cells
N2A cells
EM imaging
MERCS analysis
contacts per mitochondria
average contacts (per image)
% coverage
average % coverage (per image)
mitochondria per image
mitochondria (per image)
Control
+ATAD3A WT
+PERK WT
+ATAD3A WT +PERK WT
+ATAD3A Δ1-240
+PERK Δloop
ns
C
Control
siATAD3A
NT +Tg +Tg +GSK
ATAD3A
β-actin
ATAD3A levels
ATAD3A:actin ratio

**Supplementary Figure 3**

(A) Genotyping information for CRISPR generated PERK HA U2OS cell line. Representative western blot analysis of HA (from IP samples), PERK, eIF2α-P and eIF2α total from wild type U2OS and PERK HA U2OS cells.

(B) Schematic showing transfection of PERK or ATAD3A contructs. Graphs showing electron microscopy analysis of N2A cells expressing ATAD3A WT, PERK WT, ATAD3A + PERK, ATAD3A Δ1-240 or PERK Δloop. Graphs show: average number of individual MERCS (per image), % coverage of mitochondria by ER (per image) and number of mitochondria (per image). Statistical significance assessed using one-way ANOVA followed by Tukey's t test. For contacts per mitochondria: Control vs +ATAD3A WT (p=0.0074), Control vs +PERK WT (p=0.0431) and Control vs +ATAD3A WT +PERK WT (p=0.0212). For % coverage: Control vs +ATAD3A WT (p=0.0016), Control vs +PERK WT (p=0.0007) and Control vs +ATAD3A WT +PERK WT (p=<0.0001.) n=20 total images for each condition from 3 separate experiments.

(C) Western blot analysis showing effects of ATAD3A siRNA in N2A cells, data presented as mean ±SEM. N=3 statistical significance assessed by one-way ANOVA followed by Tukey's t test. ** denotes p=0.0002. n=3.

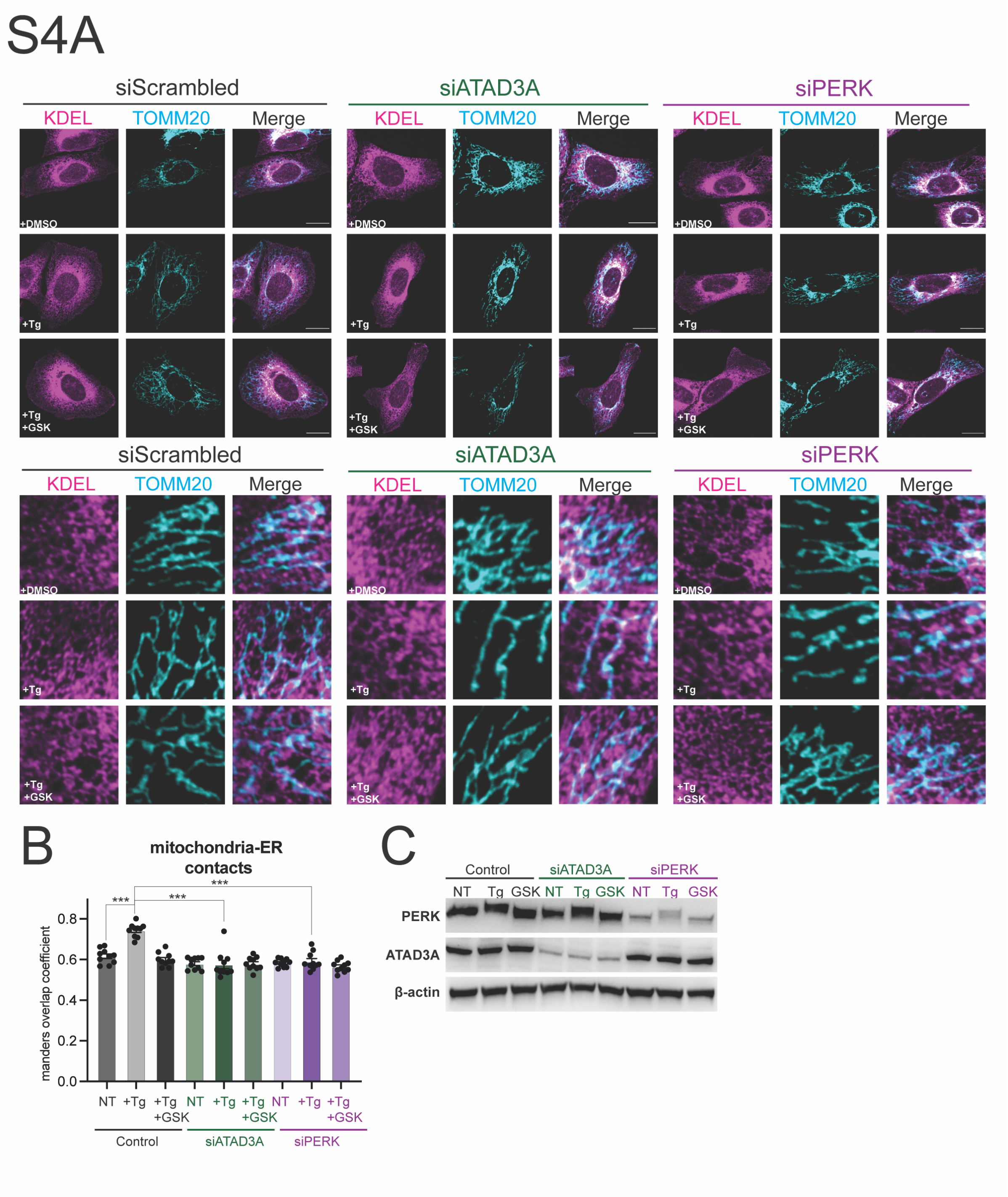
S4A
siScrambled
siATAD3A
siPERK
KDEL
TOMM20
Merge
+DMSO
+Tg
+Tg +GSK
B
mitochondria-ER contacts
***
manders overlap coefficient
0.0
0.2
0.4
0.6
0.8
NT
+Tg
+Tg +GSK
Control
siATAD3A
siPERK
C
Control
siATAD3A
siPERK
NT Tg GSK NT Tg GSK NT Tg GSK
PERK
ATAD3A
β-actin

**Supplementary Figure 4**

(A) U2OS cells were transfected with siRNAs; siControl, siATAD3a or siPERK and treated with DMSO, thapsigargin (500nM) or thapsigargin + GSK2606414 (10µM) and stained with anti-TOMM20 (mitochondria) and anti-KDEL (ER) antibodies.

(B) Colocalization of' TOMM20 and KDEL was analysed using a Mander's colocalization coefficient. Data presented as mean ±SEM. Statistical significance assessed using one-way ANOVA followed by Tukey's t test. For Control NT vs Control Tg (p=<0.0001), for Control Tg vs siATAD3A Tg (p=<0.0001), for Control Tg vs siPERK Tg (p=<0.0001) n=30 with 10 cells analysed from 3 separate experiments.

(C) Western blot analysis showing effects of PERK siRNA in U2OS cells, n=3.

**Supplementary Table 1**

Raw TMT mass spectrometry data from N2A cells treated with scrambled or ATAD3A siRNA followed by 8 hours treatment with thapsigargin (500nM) or DMSO as a control.

**Supplementary Table 2**

Normalised TMT mass spectrometry data from N2A cells treated with scrambled or ATAD3A siRNA followed by 8 hours treatment with thapsigargin (500nM) or DMSO as a control. All conditions are normalised against siScrambled DMSO treated cells.

**Supplementary Table 3**

List of mitochondrial proteins reduced on average by 20% or more in siATAD3A thapsigargin treated N2A cells when compared to siScrambled DMSO treated cells. Normalised abundances for all conditions are presented.
